# Metrics for Public Health Perspective Surveillance of Bacterial Antibiotic Resistance in Low- and Middle-Income Countries

**DOI:** 10.1101/2020.02.10.941930

**Authors:** Olga Tosas Auguet, Rene Niehus, Hyun Soon Gweon, James A. Berkley, Joseph Waichungo, Tsi Njim, Jonathan D. Edgeworth, Rahul Batra, Kevin Chau, Jeremy Swann, Sarah A. Walker, Tim E. A. Peto, Derrick W. Crook, Sarah Lamble, Paul Turner, Ben S. Cooper, Nicole Stoesser

## Abstract

Antimicrobial resistance (AMR) is a global health threat, especially in low-/middle-income countries (LMICs), where there is limited surveillance to inform empiric antibiotic treatment guidelines. Enterobacterales are amongst the most important causes of drug-resistant bacterial infections. We developed a novel AMR surveillance approach for Enterobacterales by profiling pooled human faecal metagenomes from three sites (n=563 individuals; Cambodia, Kenya, UK) to derive a taxonomy-adjusted AMR metric (“resistance potential”) which could be used to predict the aggregate percentage of resistant invasive Enterobacterales infections within each setting. Samples were sequenced (Illumina); taxonomic and resistance gene profiling was performed using ResPipe. Data on organisms causing bacteraemia and meningitis and antibiotic susceptibility test results from 2010-2017 were collated for each site. Bayesian generalised linear models with a binomial likelihood were fitted to determine the capacity of the resistance potential to predict AMR in Enterobacterales infections in each setting. The most informative model accurately predicted the numbers of resistant infections in the target populations for 14/14 of antibiotics in the UK, 12/12 in Kenya, and 9/12 in Cambodia. Intermittent metagenomics of pooled human samples could represent a powerful pragmatic and economical approach for determining and monitoring AMR in clinical infections, especially in resource-limited settings.

## Introduction

Antimicrobial resistance (AMR) is a global health emergency^1^, and imposes a particularly large socioeconomic burden in resource-limited settings, where bacterial infections and several other drivers of AMR commonly co-occur and effective antibiotics may be unavailable or unaffordable^2^. A key pillar in AMR mitigation is the development of effective and standardised AMR surveillance, to monitor trends, inform empiric treatment guidelines, identify emerging AMR threats, and monitor the impact of interventions. There has been significant investment in surveillance capacity, such as by the UK’s Fleming Fund, and an attempt to promote standardised collection, analysis and sharing of global AMR data with an emphasis on capturing clinical and microbiological information, encapsulated in the WHO Global Antimicrobial Resistance Surveillance System (GLASS)^3^. However, limitations in implementing GLASS include the time taken to develop robust infrastructural capacity to support data collection in regions where AMR is most relevant or prevalent, and the difficulty in obtaining systematic datasets even from enrolled countries with adequate infrastructure, especially outside tertiary or University centres. Surveillance strategies which could bridge or complement the implementation of approaches such as GLASS would be helpful.

Colonisation with specific species and/or drug-resistant organisms, such as nasal colonisation with *Staphylococcus aureus*^4^, or rectal colonisation with carbapenemase-producing Enterobacterales^5^, is associated with risk of infection by these organisms. Metagenomic approaches are less biased than targeted approaches which capture specific organism/resistance phenotypes of interest, and obviate the need for culturing individual organisms. Resistance gene abundances and taxonomic distributions in metagenomes are increasingly mined for a range of applications in the study of AMR, including as correlates for national antibiotic exposures^6,7^ in the case of human gut metagenomes, or as an approach to monitoring global AMR in the case of sewage^8^. However, to our knowledge, no study to date has used taxonomic and resistome profiles of pooled metagenomes to directly estimate the AMR prevalence in clinical isolates within the same population, across a range of species and antimicrobial classes. This approach would enable intermittent, strategic sampling of a subset of individuals in a population to estimate the burden of AMR in clinical isolates, facilitating evidence-based development of empiric treatment guidelines without the need for isolate-based microbiological surveillance. Most samples taken to assay colonisation (e.g. faeces/rectal swabs, nasal/throat swabs) are relatively non-invasive and acceptable for individuals, and tolerated by particularly vulnerable groups, such as neonates.

The concept of a taxonomy-adjusted AMR metric or AMR resistance potential for a metagenome has been described previously^6,9^ as the average metagenome fraction encoding resistance genes for a particular antibiotic or antibiotic group, across all bacteria in a sample that can potentially carry such resistance genes, based on known taxonomic ranges for the resistance gene families. To model the benefit of such a metric in predicting resistance in clinical isolates within a population, we took pooled faecal samples from a sub-population of individuals (>100) in three disparate geographic settings with varying AMR prevalence, namely Cambodia, Kenya and the United Kingdom (UK), and validated the model predictions using microbiological data from clinical isolates processed by laboratories in these locations over a seven-year period (2010-2017).

## Materials and Methods

### Samples and Settings

Faecal material stored in three existing biobanks was chosen for study; ethical approval for the broader use of these samples was in place. Samples comprised: (i) rectal swabs from children aged 1-59 months with and without malnutrition, taken on admission to Kilifi County Hospital in Kilifi, Kenya, from 1^st^ April to 30^th^ September 2016, and stored in Amies transport media + 1ml phosphate buffered saline at −80°C (“Pharmacokinetics of Antimicrobials and Carriage of Antimicrobial Resistance amongst Hospitalised Children with Severe Acute Malnutrition (FLACSAM)’ study^10^ [KEMRI/SERU/CGMR-C/023/3161; OXTREC 47-15]); (ii) faecal samples taken from newborns on admission to Angkor Hospital for Children in Siem Reap, Cambodia, from 11^th^ September 2013 to 10^th^ September 2014, and stored in tryptone soya broth + 10% glycerol at −80°C^11^(OxTREC ref 1047-13; this collection also included longitudinal samples taken from a subset of newborns during their inpatient stay for another study); and (iii), rectal swabs (Eswab, Copan diagnostics, Murrieta, CA, USA); 1ml Amies transport media) from individuals aged ≥18 years attending pre-admission clinics or on admission to Guy’s and St Thomas’ NHS Foundation Trust, London, UK, between February and May 2015, and stored at −80°C^12–14^ ([REC: 14/LO/2085]). Rectal swabs and faecal samples have both been used as approaches for surveying intestinal microbiota^15,16^, and are thought to give similar results^17^.

For each study site, metadata associated with microbiology tests performed on blood and cerebrospinal fluid samples (as most robustly representative of true causative pathogens) collected within 0-72 hours of admission from 01/Jan/2010-31/May/2017 were collated. Each site has a microbiology laboratory participating in external quality assurance schemes (e.g. UK National External Quality Assessment Service, NEQAS) and is additionally accredited to UK ISO15189 (London laboratory) or WHO Good Clinical Laboratory Practice standards (Kilifi laboratory). Catchment areas served by each laboratory vary: For Cambodia about two-thirds of the patients come from within Siem Reap province^18,19^; in Kenya the population served is mostly rural, within the coastal Kilifi District^20^; and in London the laboratory largely serves a South London community of approximately 0.5 million people and also regularly provides services to international patients and patients from other sites in the UK^21^. Collated metadata included bacterial species identification results, available antibiotic susceptibility testing (AST) results, specimen type and basic patient details to validate aggregate-level stratification by age. Samples were processed using standard operating procedures in accordance with accredited guidelines. In the UK, the VITEK system (bioMérieux, Marcy-l’Etoile, France) was used for AST and performed according to the British Society for Antimicrobial Chemotherapy standards^22^ (BSAC). In Cambodia and Kenya, AST was performed using a standardised disk diffusion method following the Clinical and Laboratory Standards Institute (CLSI) guidelines^23^. Where accurate AST results could not be achieved by simple disk diffusion, minimum inhibitory concentrations (MICs) were determined by Etest in both settings. The infection metadata was collated for infants < 90 days of age in Cambodia, ≤ 60 months of age in Kenya and ≥ 18 years of age in the UK.

### DNA Extraction

Samples from Cambodia and Kenya were shipped to the Nuffield Department of Medicine (University of Oxford, UK) for extraction; extractions for London samples took place at the Centre for Clinical Infection and Diagnostics Research (CIDR-King’s College London). DNA was extracted from each sample using the MoBio PowerSoil® DNA isolation kit (Qiagen, Hilden, Germany), as per the manufacturer’s instructions with optimisation steps to achieve sufficient DNA yields for sequencing (ideally ≥300ng DNA/34ul, with a view to obtaining ≥20Gbp (Giga base pairs) of data per sample). See Supplementary Methods 1 & 2. Known copy numbers of internal standards consisting of *Thermus thermophilus* HB8 genomic DNA^24^ (not normally present in faecal samples) were added to each sample prior to the addition of Solution C1 (i.e. 8.75 ul per sample [1ng/ul of Thermus DNA]). The presence of *T. thermophilus* was ascertained following sequencing by mapping reads to the Thermus reference genome.

### Sample Pooling

DNA extracts were stored at −20°C and then pooled and sequenced at the Wellcome Trust Centre for Human Genetics, Oxford, UK. For each study site, we created a “population pool”, which consisted of the pooling of equimolar concentrations of all extracts from that setting with ≥1ng DNA/μl. To validate our pooling approach, we also created one smaller pool in each setting, a so-called “30-sample pool”, which consisted of equimolar concentrations of 30 randomly selected extracts with ≥300ng DNA/34μl. An aliquot from each extract included in 30-sample pools was in turn sequenced individually for the validation study (i.e. sequenced extracts from 90 individuals in total). An aliquot from all extracts sequenced individually and included in the 30-sample pools was also included in population pools.

### Metagenomic Sequencing

Sequencing of all samples (pools and individual extracts) was performed using the HiSeq 4000 Illumina platform, generating 150bp paired-end reads (i.e. 96 metagenomes [n=90 individual metagenomes, n=3 30-sample pools, n=3 population pools]). 500ng of DNA from each sample was used for library preparation. Libraries were constructed using the NEBNext Ultra DNA Sample Prep Master Mix Kit (NEB) with minor modifications and a custom automated protocol on a Biomek FX (Beckman)^25^. At the time of sequencing, the HiSeq 4000 produced on average 72-90 Gbp of data per lane. We sequenced four individual extracts per lane to obtain on average 18-22.5 Gbp of data per sample. For the pooled samples, we sequenced one 30-sample-pool plus one population-pool per lane to obtain on average 36-45 Gbp of data per pool. Metagenomic data was obtained once for each distinct sample or pool; there were no technical replicates due to the expense of high-throughput sequencing.

### Sequence Data Processing

We determined the taxonomic abundance of bacterial species and resistance genes at individual and pooled sample levels using a recently developed bioinformatics pipeline^26^. This pipeline incorporated established approaches to taxonomic profiling, and an adapted approach to quantify resistance gene markers present in a metagenome (for details of the method, see^26^). Briefly, the sequenced paired-end reads were quality-filtered based on PHRED scores (≥ Q25 and ≥ 50 bp), and adapters removed using TrimGalore^27^. For profiling the abundance of bacterial species, the quality-filtered sequences were classified with Kraken2^28^ (v.2.0.8-beta) against bacteria, plasmid, viral and human genome sequences recovered (12 July 2019) from NCBI. With the taxonomic classification from Kraken2 and information about species specific versus non-specific genetic regions we estimated true abundance at the species level using Bracken^29^ (v.2.5.0), which was subsequently used for deriving total aggregate counts of bacterial taxa. For profiling resistance genes, the quality-filtered sequences were mapped against the Comprehensive Antibiotic Resistance Database^30,31^ (CARD, v.3.0.3) using BBMAP^32^ (v.37.72) at 100% sequence identity. The number of sequences that mapped to each resistance gene were subsequently corrected to remove resistance gene length bias. This was done using four metrics, namely (1) specific read count (number of sequences that map exclusively to the resistance gene); (2) specific lateral coverage (proportion of the resistance gene covered by sequences mapping exclusively to the gene); (3) resistance gene length; and (4) and average read length (average length of reads that mapped to the resistance gene), and by the following formula: corrected gene count (CGC) = (specific read count x average read length) / (resistance gene length x specific lateral coverage).

The CARD database attempts to classify each resistance gene variant by its association with AMR. To be included in CARD, an AMR determinant must be described in a peer-reviewed scientific publication, have its DNA sequence available in GenBank, and include clear experimental evidence of elevated MIC over controls^31^. We used these data to map and aggregate counts of resistance genes/variants associated with resistance to a specific antibiotic. In the process, we ranked the resistance genes/variants into two categories, reflecting to some extent the public health risks posed^33^, and thereby creating two sets of antibiotic resistance gene metrics. The first (AMR_DEF_; Supplementary Data 1), included only AMR determinants with the “*Confers_Resistance_to_Antibiotic*” relationship ontology term, whereby the gene associated with demonstrably elevated MIC is known to confer or contribute to clinically relevant resistance to a specific antibiotic drug^31^. The second (AMR_ALL_; Supplementary Data 2), contained corrected counts of all resistance genes with clear experimental evidence of increasing the MIC, including those associated with clinically relevant resistance (as for AMR_DEF_), plus those without the “*Confers_Resistance_to_Antibiotic*” relationship ontology term. For the purposes of this study we have used the term “resistance gene” to define any relevant genetic marker of resistance, including genes that confer resistance by mutation (but can have a susceptible wild type), and genes that confer resistance through presence/absence.

### Validation of Pooling

We evaluated to what extent pooled resistome data was a non-biased representation of the individual resistomes making up the pool. Resistance gene abundances of the 30-sample pools and individually sequenced samples were converted to relative abundances, such that gene abundances in each sample summed to one. Then, for each of the three different settings, individual samples were used to compute the empirical distribution of each gene by repeated random sampling of its relative gene abundance out of the individual samples (bootstrapping with *n*=100,000 repeats). We were then able to compare the pool abundance of each gene with its empirical distribution in the same setting (within-setting comparison) and in the other two settings (across-setting comparison). We computed the fraction of resistance genes for which the pool estimate was within 90% central quantile of the empirical distribution. The resulting metric was restricted between 0 (i.e. 0% of resistance genes in the pool were as expected given the individual resistomes) and 1 (i.e. 100% of resistance genes in the pool were as expected). Because bootstrapping of gene abundances relies on having a sufficient number of samples with non-zero abundance, we limited our analysis to genes present in ≥50% of all individual samples (n=121 genes). Given the central quantile choice above (i.e. 90%), a value of ∼0.90 would imply a non-biased representation of individual resistomes by the pooled resistome. For visualization, non-metric multidimensional scaling, an ordination-based method, was used to show pair-wise dissimilarities between resistomes from population pools, 30-sample-pools and individual sample means within and across settings. Individual sample means for each setting were, for each AMR gene, the sum of CGCs across all individually sequenced samples.

### Taxonomy Adjusted Resistance Potential Metrics

We developed several candidate metrics of resistant infection risk, based on pool metagenomic data on resistance gene abundance and bacterial species composition, and evaluated their potential to accurately predict the likelihood of antibiotic resistant invasive infections in a population. We refer to these as ‘taxonomy-adjusted resistance potential (RP)’ metrics, which consisted of two parameters. The first parameter, *R*_*CGC*,_ was given through the sum of corrected gene counts (CGC) of variants associated with resistance to a given antibiotic, *j (R*_*CGCj*_) divided by the total CGC of all resistance genes in the pool. *R*_*CGC,j*_ was calculated based on either variants with experimental evidence of increasing the MIC (AMR_ALL_) or only variants known to confer clinically relevant resistance (AMR_DEF_). The second parameter, *R*_*Tax*_, was given through the estimated abundance of a clinically relevant bacterial grouping (derived from Bracken estimates) divided by the total estimated abundance of bacterial taxa in the pool. The bacterial groupings tested were the Enterobacterales order, Enterobacteriaceae family, and the grouping of the four most common and clinically relevant bacterial genera/species within the Enterobacteriaceae family across sites (namely *Escherichia coli, Klebsiella pneumoniae, Salmonella* spp, *Enterobacter* spp).

### Bayesian Modelling

With each taxonomy-adjusted RP, we fitted a Bayesian generalized linear model to the data and applied model comparison. This allowed us to assess the potential of the different metrics to predict observed antibiotic resistance amongst clinical invasive Enterobacterales isolates. We used de-duplicated counts of isolates (unique bacterial species per antibiogram and patient-ID) for the analyses. We let *i* denote the setting (Cambodia, Kenya or UK), and *j* the antibiotic (see below for a list). We assumed that counts of resistant samples follow a binomial distribution. Our model then predicts the count of resistance (*r*_*i,j*_) among tested Enterobacterales isolates (*n*_*i,j*_) using a probability of resistance (*p*_*i,j*_), which is modelled as

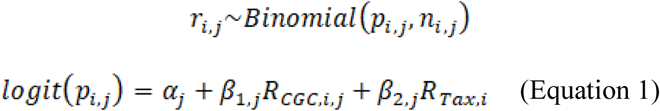

The model intercept (*α*) is specific for each antibiotic (*j*) but not setting (*i*), representing a baseline propensity of resistance for any given antibiotic. Because resistance propensities can vary widely between different antibiotics. We assume independent baselines (fixed effects). The setting-specific information is *R*_*Tax,i*_, which gives information about pathogen levels in setting *i*, as well as *R*_*CGC,i,j*_, which carries information about resistance toward antibiotic *j* in setting *i*. For *β*_1,j_ and *β*_2,j_, the predictive effects of *R*_*CGC*_ and *R*_*Tax*_, we assumed these to represent the clinical ecology of resistance genes so that they are specific to each antibiotic, *j*, but not to each setting, *i*. We further assumed that different antibiotics have different but related *β*-values (variable effects, specified below). We included only those antibiotics that had existing antibiotic susceptibility test (AST) data in at least two out of three settings (trimethoprim-sulfamethoxazole, nitrofurantoin, nalidixic acid, meropenem, imipenem, gentamicin, ciprofloxacin, chloramphenicol, cefuroxime, ceftriaxone, ceftazidime, cefpodoxime, cefoxitin, cefotaxime, ampicillin, amikacin); missing observations were excluded from the likelihood evaluation. We fitted the above model with *R*_*CGC*_ being either AMR_DEF_ or AMR_ALL_ and with *R*_*Tax*_ being either of the three bacterial groupings discussed earlier, yielding a total of six separate model fits. Due to the limited number of infection isolates with AST results (especially in Cambodia), we chose standard weakly informative priors for the intercept and the effect parameters. In addition, we restricted the effect of gene abundance to be positive, reflecting our view that only a positive association of resistance genes and clinical resistance is biologically reasonable. We therefore chose

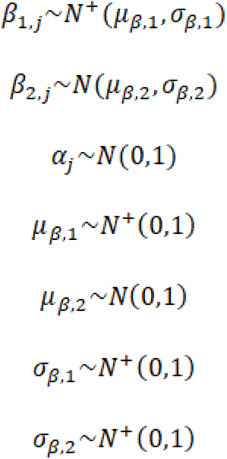

where *N* denotes a normal distribution and *N*^+^ denotes a half-normal distribution covering only positive values. Each model was fit using Stan software^34^ (v2.19.1), with which we sampled 50,000 samples after a burn-in period of 5,000 samples using four independent chains.

The best taxonomy-adjusted RP metric was selected using Bayesian leave-one-out cross validation^35^ which estimates a model’s pointwise out of sample prediction accuracy. The prediction accuracies are then used to directly compare all models using stacking weights^36^. In brief, models with smaller cross-validation errors (e.g. smaller prediction errors), get more weight relative to other models in the model comparison. We also included in the comparison two models with *R*_*CGC*_ (either AMR_DEF_ or AMR_ALL_), following Equation (1), but without *R*_*Tax*_. Finally, the overall value of using any taxonomy-adjusted RP metric for predicting clinical resistance was assessed by including in the model comparison a baseline model without predictors. The prediction accuracy of taxonomy-adjusted RP was also assessed visually by comparing the best model’s predictions of sample counts of resistance (and their 95% credible intervals [CI]) against the observed counts (Figure 5). For settings and antibiotics where zero samples were tested, we imputed the sample size by computing the rounded mean of the sample sizes of the other two settings. Model comparisons and all further data analyses were performed in R-3.6.1 statistical software^37^. The dataset used for the Bayesian modelling is given in Supplementary Data 3.

**Fig 1.**
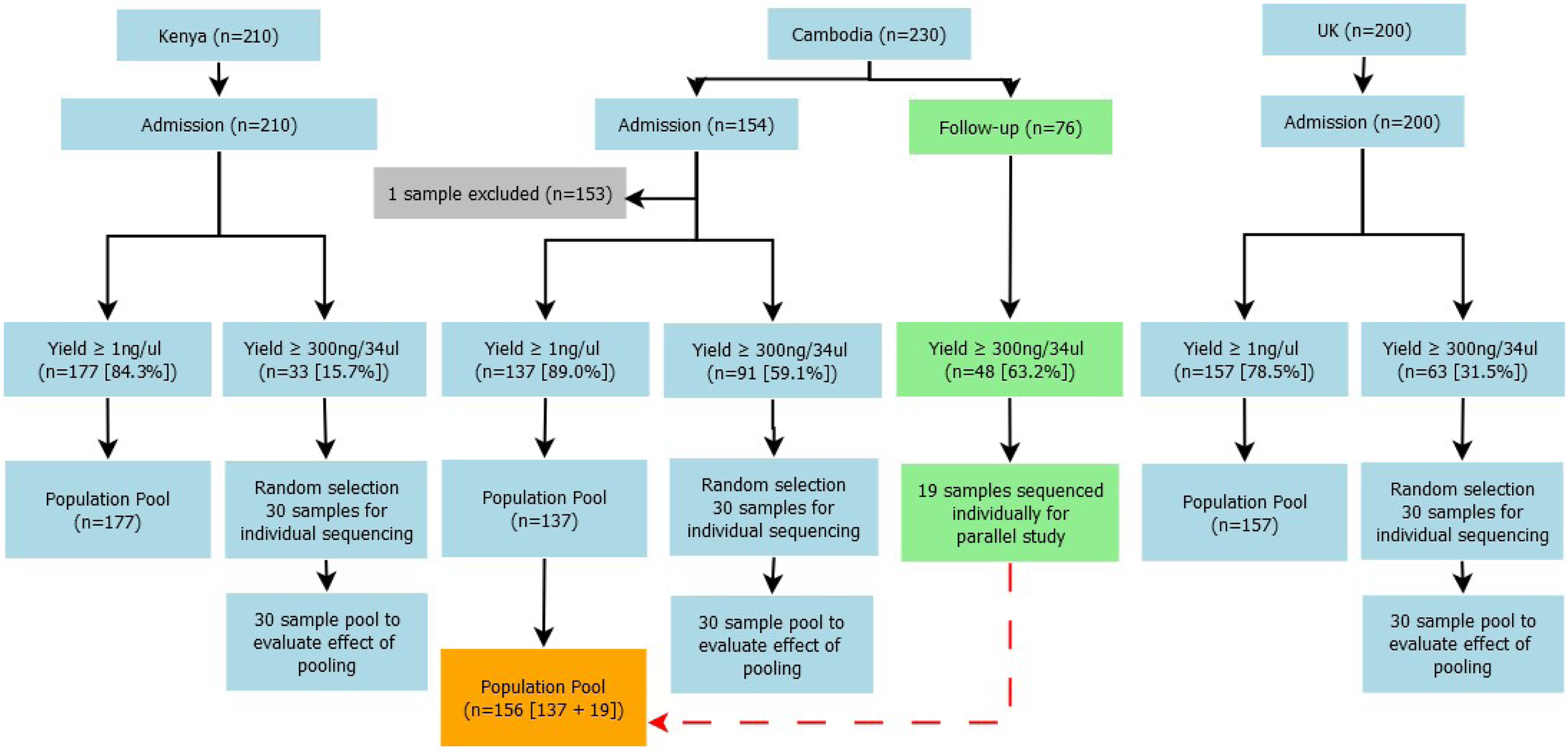
Sample Processing Workflow. We identified 863 different antimicrobial resistance genes across any sample or pool (Cambodia: 684; Kenya = 527; UK = 520), which were proven to increase the MIC for 163 antimicrobials (AMR_ALL_) and known to confer clinically relevant resistance for 113 antimicrobials (AMR_DEF_). The number of resistance genes identified in population pools was largest in Cambodia (n=490), followed by the UK (n=389) and Kenya (n=386). The median number of resistance gene types identified per individual sample was also higher in Cambodia (median=162; IQR= 126-187 [Min-Max =33-231]), followed by in Kenya (median=143; IQR= 127-205 [Min-Max =97-256]) and UK (median=134; IQR= 126-148; [Min-Max =61-217]).

**Fig 2.**
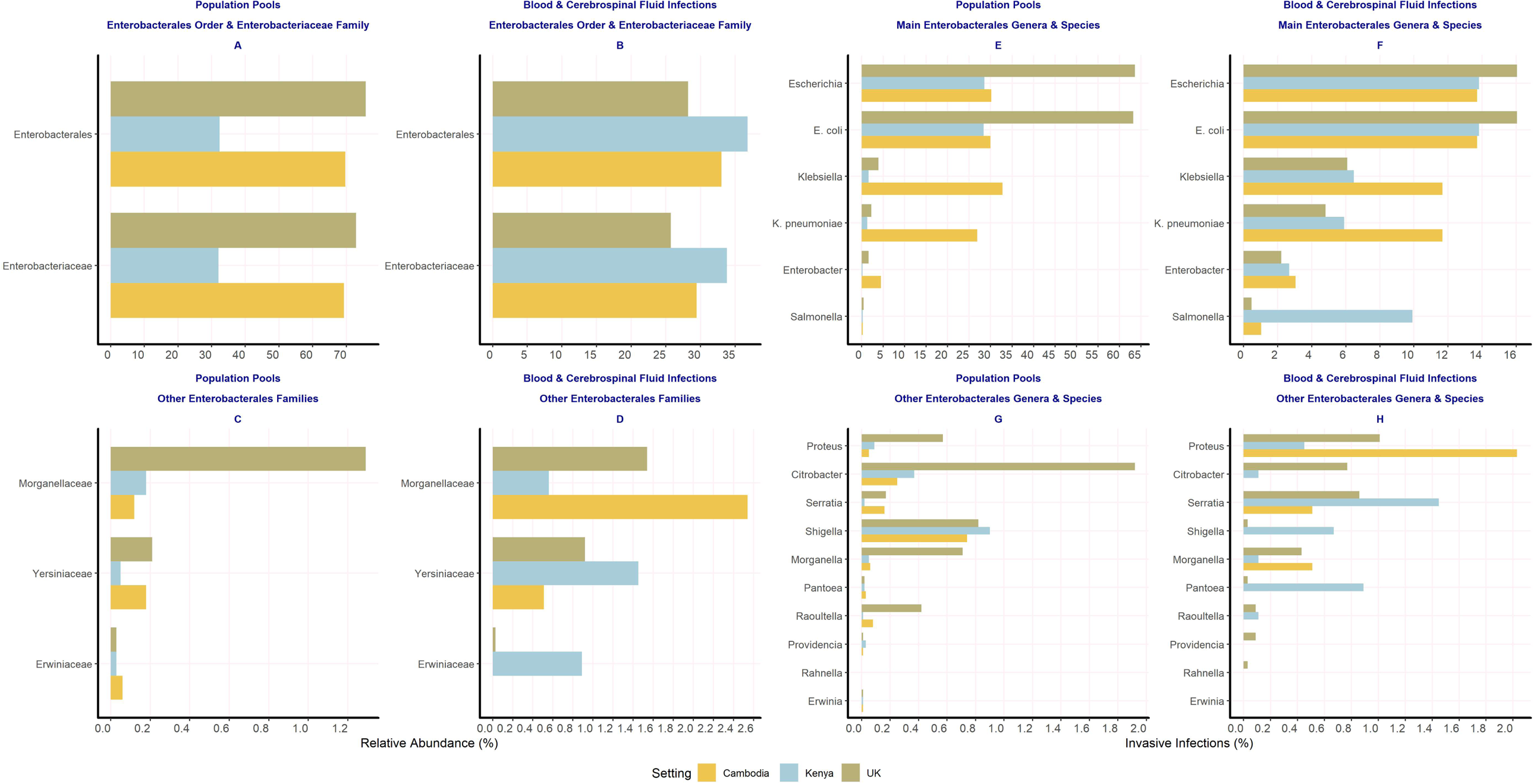
Bacterial (Enterobacterales) taxa in population pools and in blood and cerebrospinal fluid infections from Cambodia, Kenya and UK. Panels for population pools (A, C, E, G) show, for each setting, the abundances of Enterobacterales taxa divided by the total abundance of bacterial taxa in a pool. Abundances are derived from Bracken estimates. Panels for invasive infection data (B, D, F, H), show percentages of Enterobacterales infection isolates out of all bacterial infection isolates with speciation results identified from blood and cerebrospinal fluid samples in target age groups, in each setting, from 2010-2017 (Cambodia [n=197]; Kenya [n=910]; UK [n=3356]).

**Fig 3.**
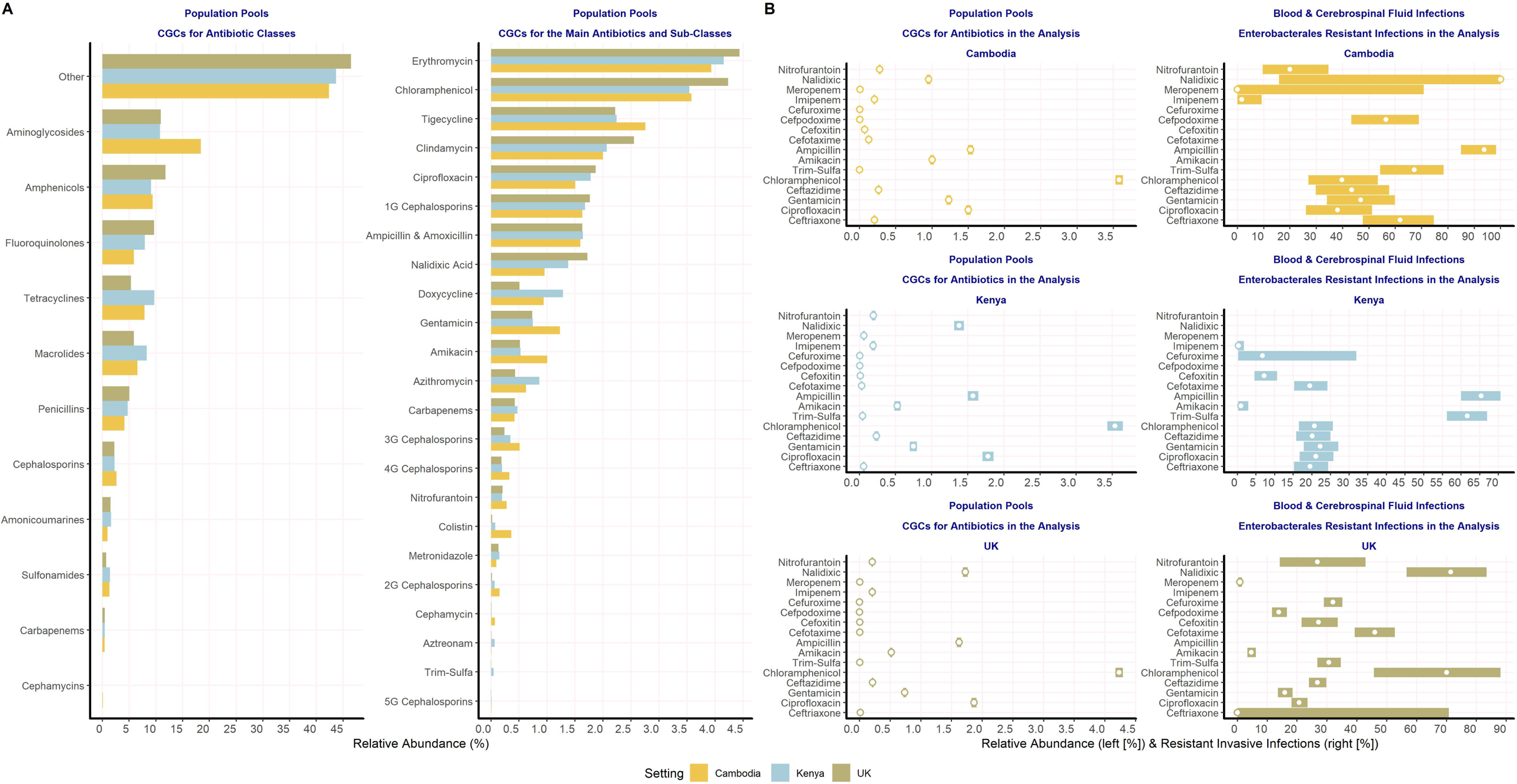
Relative abundance of resistance gene counts in population pools and percentage of resistant Enterobacterales blood and cerebrospinal fluid infections in Cambodia, Kenya and UK. Panels in Fig 3A show, for each setting, corrected resistance gene counts (CGCs) for a given antibiotic, antibiotic class, or sub-class, divided by the total corrected AMR gene counts identified in the population pool. Relative abundances were calculated using AMR_ALL_, which considers corrected counts of genes and variants (CGC) increasing the MIC or conferring clinically relevant resistance for a given antibiotic. Panels in Fig 3B show, for each setting, the observed percentage of Enterobacterales resistant infections for 16 antibiotics with AST data in ≥ 2 settings (right-hand side), and the relative abundance of CGCs for the same antibiotics in population pools, based on AMR_ALL_ (left-hand side). Percentages are shown with 95% exact binomial confidence intervals in both panels.

**Fig 4.**
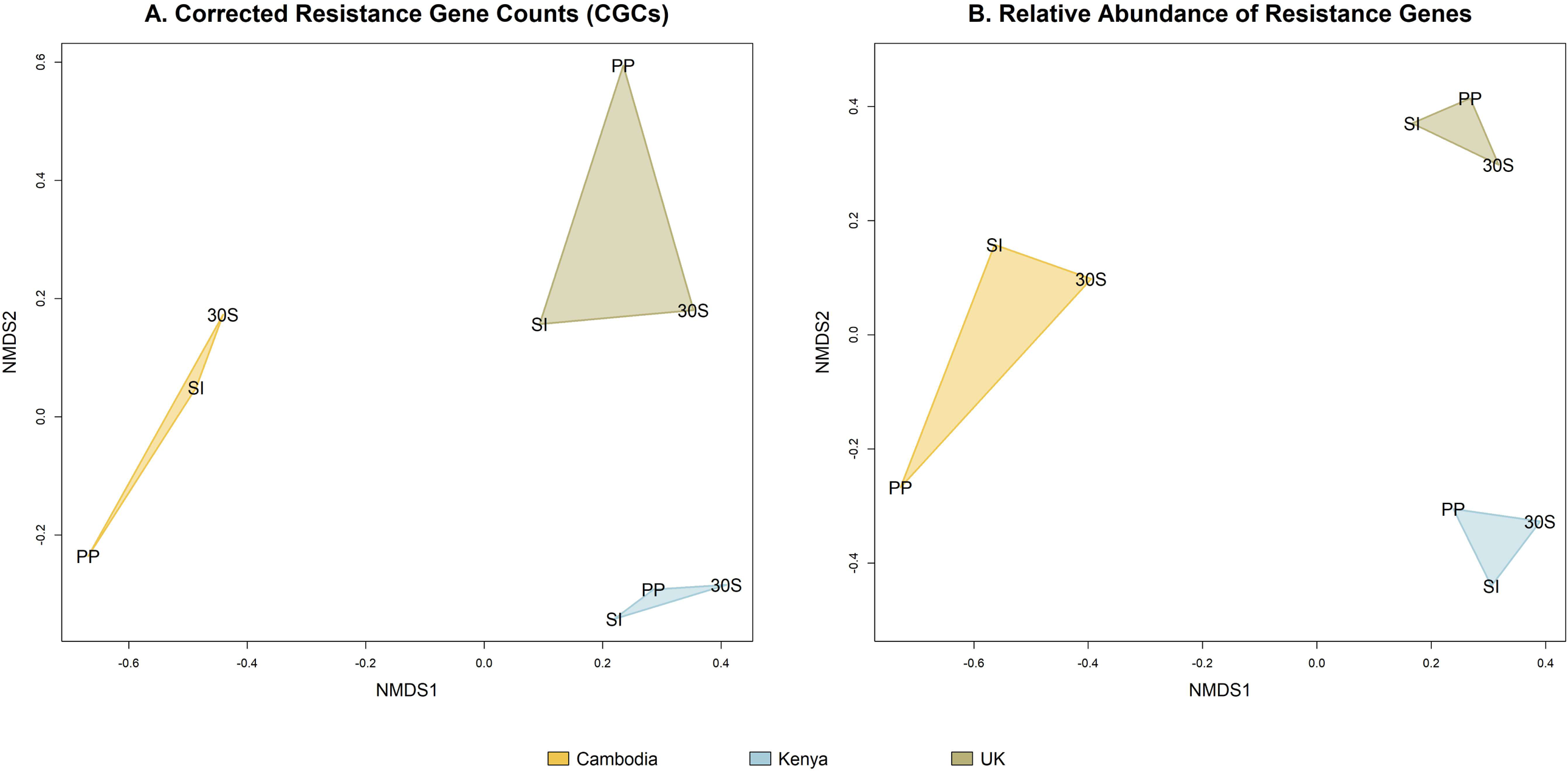
Pair-wise dissimilarities in the resistome of population pools, 30-sample-pools and individual sample means within and across settings. Using non-metric multidimensional scaling (NMDS) ordination-based method, Fig 4 shows pair-wise dissimilarities of resistance gene counts from population pools (PP), 30-sample-pools (30S) and individual sample means (SI) within and between settings, following mapping of sequences from individual and pool metagenomes against CARD and a correction to remove resistance gene length bias from counts. Dissimilarities are shown for the absolute corrected resistance gene counts (CGC; left hand-side) and the relative abundance of resistance genes (right hand-side). Relative abundances for genes in pools were calculated by dividing the CGC for each gene by the total CGC of all resistance genes in the pool. Individual sample means were, for each resistance gene, the sum of CGC across all individually sequenced samples. This, divided by the total CGC of all resistance genes across all individually sequenced samples, was the relative abundance of each resistance gene based on individual sample means.

**Fig 5.**
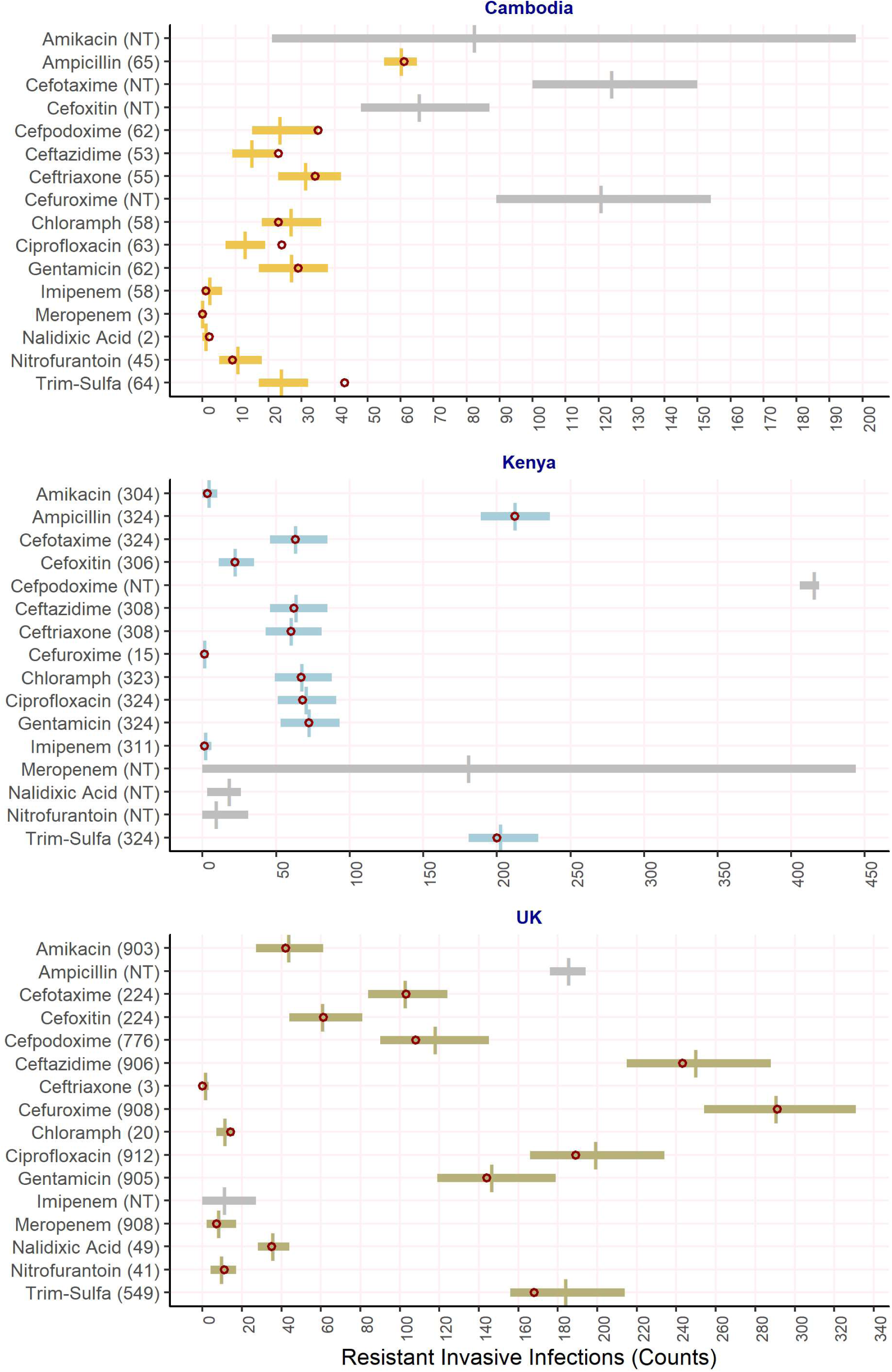
Bayesian model prediction of numbers of Enterobacterales invasive infections with resistance to antibiotics with antibiotic susceptibility test (AST) results in ≥2 settings. Horizontal bars represent 95% highest density posterior interval and vertical lines represent means of the predicted resistant sample counts based on the model using metagenomic data from population pools. Coloured bars (yellow: Cambodia; blue: Kenya; brown: UK) are shown where clinical data on resistance (i.e. AST) was available and grey bars where it was not. For grey bars the sample size was imputed. Red circles show the number of blood and cerebrospinal fluid Enterobacterales infections that were found to be resistant to the antibiotic listed in the y-axis. The number of isolates with AST results are also given in the y-axis. The red circle is missing where no AST results were available. In cases where there is minimal uncertainty in the model estimate, the red circle may overshadow the 95% credible interval bars (e.g. meropenem [Cambodia]; cefuroxime [Kenya]). “Trimethoprim.” is short for trimethoprim-sulfamethoxazole; “Cloramph” is short for chloramphenicol. NT = no AST data available.

## Results

The study included 210 admission samples from Kenya, 200 from the UK and 153 from Cambodia (n=154 – 1 rejected sample), totalling 563 samples for metagenomic analysis (Fig 1). In addition, 76 follow-up samples were taken from 37/154 newborns in Cambodia during their inpatient stay or upon hospital discharge for a separate project; these were processed alongside the study samples (Fig 1). We only considered DNA extracts with yields ≥1ng/ul (79-89% of samples; Fig 1), and 19 DNA extracts from the separate longitudinal study were included in the Cambodia population pool due to processing error. In total, population pools in Kenya, the UK and Cambodia, comprised 177, 157, and 156 pooled sample extracts. Thirty high DNA-yield samples (≥9ng/ul) from each setting were used for the validation study, as well as being included in the population pools. To prevent bias, potential associations between high-yield samples and population traits were ruled out in advance. The total Gbp of data obtained per population pool were 51.6 (Kenya), 55.1 (UK) and 52.6 (Cambodia). The median Gbp obtained for individually sequenced samples were 24.2 (Kenya), 22.1 (Cambodia), 22.4 (UK).

A summary of the Enterobacterales taxa identified from population pools and invasive infections in each setting is given in Fig 2. Enterobacterales were the main bacterial taxa identified from population pools in the UK (75.7%) and Cambodia (69.7%) but not in Kenya (32.4%) (Fig 2A). Within the Enterobacterales, >95% of the bacteria were from the Enterobacteriaceae family in all settings (UK: 96.3%; Cambodia: 99.4%; Kenya: 99.1%). The predominant species within the Enterobacterales order in population pools were *E. coli* and *K. pneumoniae*, followed by *Enterobacter* spp. (Fig 2E). These species and genera combined accounted for 92.4% of all Enterobacterales taxa in Kenya, 88.5% in the UK and 88.1% in Cambodia. The abundance of *E. coli*, was >20-fold higher than that of *K. pneumoniae* in population pools from the UK (*E. coli*: 63.2%; *K. pneumoniae*: 2.2%) and Kenya (*E. coli*: 28.4%; *K. pneumoniae*: 1.3%). In contrast both species had similar abundance in the Cambodia population pool (*E. coli*: 30%; *K. pneumoniae*: 26.9%). *Enterobacter* spp. abundance was also higher in Cambodia (4.5%) compared to the UK (1.6%) or Kenya (0.2%). The remaining Enterobacterales comprised other genera, each being <2% of the total bacterial taxa in the three settings (Fig 2G). Infections by Enterobacterales accounted for approximately a third of all blood and cerebrospinal fluid infections in the three settings (Kenya: 36.8%; Cambodia: 33.0%; UK: 28.2%) (Fig 2B). Similar to the findings from population pools, most of these Enterobacterales infections involved the Enterobacteriaceae family (UK: 91.2%, Cambodia: 89.2%; Kenya: 91.8%; Fig 2B). Likewise, the predominant Enterobacterales species in all settings were *E. coli* and *K. pneumoniae*, with the proportion of *E. coli* infections being at least double that of *K. pneumoniae* in the UK (*E. coli*: 16.1%; *K. pneumoniae*: 4.8%) and Kenya (*E. coli*: 13.8%; *K. pneumoniae*: 5.9%), but not in Cambodia (*E. coli*: 13.7%; *K. pneumoniae*: 11.7%) (Fig 2F). *Enterobacter* spp. was the next most common Enterobacterales genus in all settings (Cambodia: 3.1%; Kenya: 2.7%; UK: 2.2%), but the remaining Enterobacterales species and genera accounted for <2% of the total invasive infections by any bacterial order each in all three settings (Fig 2H). A notable exception was *Salmonella* spp., which accounted for 9.9% of the total infections in Kenya (therefore also included in Fig 2F and the equivalent plot for population pools [Fig 2E]). Details of all invasive infections by bacteria other than the Enterobacterales are given in Supplementary Fig 1.

The highest relative abundances of resistance genes observed in each setting were for genes associated with resistance to aminoglycosides, amphenicols, fluoroquinolones, tetracyclines and macrolides (48.1%, 45.8% and 43.6% of the total counts in Cambodia, Kenya and the UK respectively) (Fig 3A, left-hand panel). Relative abundance of resistance genes associated with these five broad antibiotic classes differed between settings. For example, the relative abundance of resistance genes for aminoglycosides in Cambodia (18.4%) was almost double that in Kenya (10.8%) or UK (10.9%). The next highest relative abundance was of genes conferring resistance to penicillins (Cambodia: 4.1%; Kenya: 4.7%; UK: 5.0%) and cephalosporins (Cambodia: 2.6%; Kenya: 2.3%; UK: 2.2%). Resistance gene counts for other antibiotic classes were <2% of the total gene counts in all settings, including to carbapenems (Kenya [0.5%], Cambodia and UK [0.4%]). For single antibiotics or antibiotic sub-classes (e.g. 1^st^ generation cephalosporins), the highest relative abundances were observed for erythromycin (Cambodia: 3.9%; Kenya: 4.2%; UK: 4.4%) and chloramphenicol (Cambodia: 3.6%; Kenya: 3.5%; UK: 4.2%) in all settings (Fig 3A, right-hand panel). That for resistance genes to antibiotics other than those listed was 76% (Cambodia), 76.5% (Kenya) and 76.1% (UK) (data not shown). The relative abundance of resistance genes for all other single antibiotics/antibiotic sub-classes was <2% in all settings, except for tigecycline (Cambodia: 2.8%; Kenya and UK: 2.2%) and clindamycin (Kenya: 2.1%; UK: 2.6%). Resistance prevalence in Enterobacterales isolates causing blood and cerebrospinal fluid infections is displayed in Fig.3B (right-hand panel) for the 16 antibiotics with antibiotic susceptibility test data in ≥2 settings. For comparison, this is shown alongside the relative abundance of resistance genes for the same antibiotics in population pools (Fig.3B, left-hand panel).

Pair-wise dissimilarities in resistomes from population pools, 30-sample-pools and individual sample means (i.e. sum of CGC for the resistance gene types across all individually sequenced samples) were calculated both within and across settings (Fig 4A and 4B), considering either the absolute CGC values for each resistance gene type or their relative abundance based on the CGC values. Population pools, 30-sample pools and individual sample means were less dissimilar and hence more closely related within settings than across settings. In addition, within each setting, individual sample means were more often less dissimilar to 30-sample pools than to population pools. In Cambodia 362 AMR genes were identified in the 30-sample pool compared to 616 across all 30 individual samples. The 30-sample pool in Kenya comprised 339 genes compared to 499 across all individual samples. Finally, in the UK 318 AMR genes were identified from the 30-sample pool compared to 422 across all individual samples. However, when comparing individual samples and pools from the same setting quantitatively, the average fraction of resistance genes for which the 30-sample pool estimate was within the central interval of the empirical distribution inferred from individually sequenced samples was 97% (Kenya: 98%; Cambodia: 97%; UK: 95%). In contrast, the average fraction was 86% across comparisons between different settings (min-max: 80-92%). All 30-sample pool resistomes therefore had substantially higher similarity to individual resistomes from the same setting relative to the comparison with other settings.

The best taxonomy-adjusted RP metric - resulting in the highest point-wise out of sample prediction accuracy and the greatest relative model weight - used the taxonomic parameter *R*_*tax*_ measuring *Escherichia coli, Klebsiella pneumoniae*, Salmonella spp. and Enterobacter spp., and the abundance of resistance genes increasing the MIC or conferring clinically relevant resistance (AMR_ALL_ version of the *R*_*CGC*_ metric). This AMR_ALL_ model outperformed the other models, including a baseline model without any metagenomics information, plus those models without taxonomic (*R*_*tax*_) information (Bayesian model averaging weights: Baseline [no *R*_*CGC*_ and no *R*_*tax*_] = 0; *R*_*CGC*_ only [No *R*_*tax*_] = 0; Best model = 0.47]. Supplementary Data 4).

Model predictions were made for 16 antibiotics, which were those that had antibiotic susceptibility test (AST) data for Enterobacterales isolates causing infection in at least two of the three settings (Supplementary Data 5). Our best model accurately predicted the number of resistant infections in the target populations for 100% of antibiotics with AST data in Kenya (12/12) and UK (14/14). In Cambodia, the model accurately predicted the counts of resistant infections for 75% of antibiotics (9/12). Compared to this, the baseline model did not correctly predict 50% of antibiotics across the three settings (19/38). We computed the mean-squared errors of the mean model predictions relative to the observations. The baseline model had an error of 468, whilst the final model (Fig 5) had an error of 33.

Bayesian model predictions expressed as percentages are shown in Supplementary Fig 2 for antibiotics where AST results were available from > 100 invasive infection isolates. Above this threshold, predicted percentage resistance was accurate for 100% of antibiotics (14/14 with >100 tested isolates).

## Discussion

In this study we have demonstrated the feasibility of a novel, pragmatic approach to surveillance of bacterial antimicrobial resistance of relevance to human infection, with a focus on Enterobacterales as one of the major bacterial resistance threats^38,39^. Our results show that metagenomic analysis of pooled faecal material (pooled at equimolar concentrations) is effective at predicting invasive infections caused by Enterobacterales resistant to in-use antibiotics in a population, across a range of different age groups and geographic settings. Our approach would enable intermittent, acceptable and relatively non-invasive sampling of a small number of individuals within a population (e.g. 100-200), with the advantage that a single centralised infrastructure (either in-country or internationally) could undertake the metagenomic sequencing and analysis. This can be done independently of development of a network of classical microbiological laboratories in multiple settings, which can be resource-intensive in terms of capital and running costs, and is not feasible in the short-term, especially in LMICs, which frequently have the highest AMR burden.

Based solely on pool size and sequencing depth (50-55Gbp/pool), we developed predictive metrics (RP) without the need for costly and labour-intensive multiplexing of samples (i.e. individually identifying samples in the pool by means of barcoded sequences) or selective sequencing approaches based on enrichment for predefined panels of resistance genes. Unlike other AMR gene profiling approaches our bioinformatics pipeline (ResPipe) incorporates the capacity to identify both specific AMR gene variants (e.g. such as *bla*_CTX-M-33_ versus *bla*_CTX-M-63_), as well as being able to aggregate by gene family. This is especially important for the prediction of phenotypes, as genes that differ by only single nucleotides/amino acids can have distinct phenotypic spectra. Pooled metagenomes/resistomes were also found to be an accurate, non-biased representation of the individual sample metagenomes/resistomes. Population pools comprising rectal swabs with as little as ≥1ng/ul DNA/sample were found to be sufficient to derive RP metrics with predictive value; this is useful in terms of optimizing the sample processing workflows. Finally, in producing relatively deeply sequenced (50-55Gbp/metagenome) and complete (i.e. not restricted to 16S) metagenomes on 90 individuals, we have also made a significant contribution to the human microbiomics data repository, freely available for other researchers to use for study.

The limitations of our approach were most obvious for the neonatal group from Cambodia, where predicted resistance matched the observed resistance in invasive isolates for 75% of antibiotics compared to 100% of antibiotics in Kenya and the UK. One explanation for this might be that the population pool for this group was found to have included 19 longitudinal samples (12% of all samples in the pool) collected from individuals during their hospital inpatient stay, potentially biasing the metagenomics profile of population pools and infection metadata designed to reflect community (i.e. non-hospital) profiles. Rapid changes in the neonatal resistome occur following exposure to the hospital environment^40^. Analyses of neonatal metagenomes have shown that these are predisposed to rapid flux, and in hospital typically reflect the environmental hospital “microbiome”^41^. Cambodia was also the only setting where the age group considered for metagenomics analysis (i.e. neonates), did not correspond exactly with the available infection metadata analysed (i.e. infants up to 90 days of age), which may also have influenced the accuracy of our predictive approach. Our analysis was also limited by the scarce antibiotic susceptibility test (AST) results available for invasive infection isolates, particularly in Cambodia, where the maximum number of isolates with AST results for any given antibiotic was 65, compared to 324 in Kenya and 912 in UK. The smaller number of isolates from Cambodia meant that the model fit contained less information to accurately predict resistance in this setting. Moreover, AST results were only available for a limited number of antibiotics across all three settings, and ideally AST approaches used for comparison would have been standardised across the settings. Finally, our analyses are heavily dependent on the robustness of the reference gene database, and the accuracy of genotypic-phenotypic correlations catalogued therein. In general, however, we would expect this knowledge base to become increasingly robust, thus strengthening our predictions. This may explain why in this study, a model that considers all gene variants with experimental evidence of increasing the minimum inhibitory concentration (MIC), outperformed a model considering only genes known to confer clinically relevant resistance.

Further studies to validate our promising proof-of-principle observations in additional settings across age categories, especially the neonatal group, are warranted. There is potential to extend the approach to consider other priority bacterial groups and different colonisation samples. For example, pools could be extended to include samples from nasopharyngeal sites, where other potential pathogens predominate (e.g. *Streptococcus* spp., *Staphylococcu*s spp.). To develop the most rapid, convenient, simple and inexpensive method possible, future studies should also consider further simplifications to the method such as whether the same accurate predictions can be generated by pooling all samples prior to DNA extraction and then performing the extraction only once. Further work should also test the resolution of the approach to characterise and track local/sub-national variation in AMR prevalence, or in community versus healthcare-associated contexts. A mathematical framework for minimum-cost implementation of pooled-sample metagenomics-based surveys to quantify the burden of resistance in new settings without prior microbiology or AST data would also be of benefit, and could be greatly informed by the data we have generated, which can contribute to simulation work addressing pools sizes, pool numbers per region, and sequencing depth.

We conclude that surveillance based on population colonisation metagenomics and taxonomy-adjusted AMR metrics presented here are in principle a valuable public health opportunity, and may represent an alternative or bridging measure to the implementation of local and regional laboratory-based infrastructures focussed on culturing isolates from clinical specimens, especially in resource-limited settings. This novel approach could be used to overcome the current paucity of quality AMR surveillance data and inform setting-tailored rationalization of/or access to antibiotics, context-appropriate treatment guidelines, organized measures to prevent AMR and ultimately public-health decision in conjunction with relevant stakeholders, especially in LMICs.

## Ethics Declarations

This research was conducted with approval from the Oxford Tropical Research Ethics Committee (OxTREC Reference: 5126-16) following local ethics clearance for inclusion of Cambodia and Kenya sample collections, and approval of a substantial amendment to 14/LO/2085 by the National Research Ethics Service (NRES London – Camberwell St Giles), for inclusion of the London sample collection in this study.

The authors declare no competing interests.

## Data and Code Availability

The raw sequence data reported in this study have been deposited in the European Nucleotide Archive under accession number PRJEB34871. The code to extract CARD data, that required to generate the final datasets and analyses, plus any required input files, are available from the ResPipe GitLab repository (https://gitlab.com/hsgweon/ResPipe); ResPipe output data can be found at the ResPipe Gitlab subdirectory (https://gitlab.com/hsgweon/ResPipe/tree/master/data).

## Author Contributions

This work was first conceived by O.T.A., with support from N.S. and B.S.C.; O.T.A, N.S., B.S.C., R.N. and H.S.G. designed the study. K.C., J.W., O.T.A. and N.S. developed and validated modified DNA extraction protocols for this study. K.C., J.W. and R.B. conducted or facilitated most of the pre-sample-pooling laboratory work. S.L. designed the methods and provided technical guidance for sample pooling and sequencing and conducted the sequencing work. J.A.B., J.D.E., P.T. and R.B. facilitated the collation and transfer of samples and data from participant settings. They also provided technical support for clinical and microbiology study procedures and for the development of context-appropriate standard operating procedures. N.S., A.S.W., T.E.P., D.W.C. and B.S.C. provided support and guidance for all technical aspects of the study (including for bioinformatics and data analyses) and contributed to the revision of study outputs. T.N. contributed to the mining, standardisation and analysis of infection metadata from each setting. H.S.G. conducted the bioinformatics work, designed the methods for corrected gene counts and extracted the data from CARD. J.S. provided the computing support for the study. O.T.A. conducted mining, linkage and visualisation of study data. R.N. conducted the validation and Bayesian analyses and B.S.C. contributed to revision of these methods. O.T.A, N.S., R.N. and H.S.G. produced the first manuscript draft. All authors contributed significantly to the iterative review of the draft.

## Acknowledgements

We are grateful to Professor Mike English (Centre for Tropical Medicine and Global Health, University of Oxford, UK), for facilitating the initiation of this work by bringing together some of the groups involved in the research. We are also grateful to Dr Alessandra Natale, for supporting the laboratory work for this study at the CIDR (Centre for Clinical Infection and Diagnostics Research, Department of Infectious Diseases, King’s College London, UK). The study was funded by Bill & Melinda Gates Foundation (grant agreement OPP1160974) and was sponsored by University of Oxford.

---

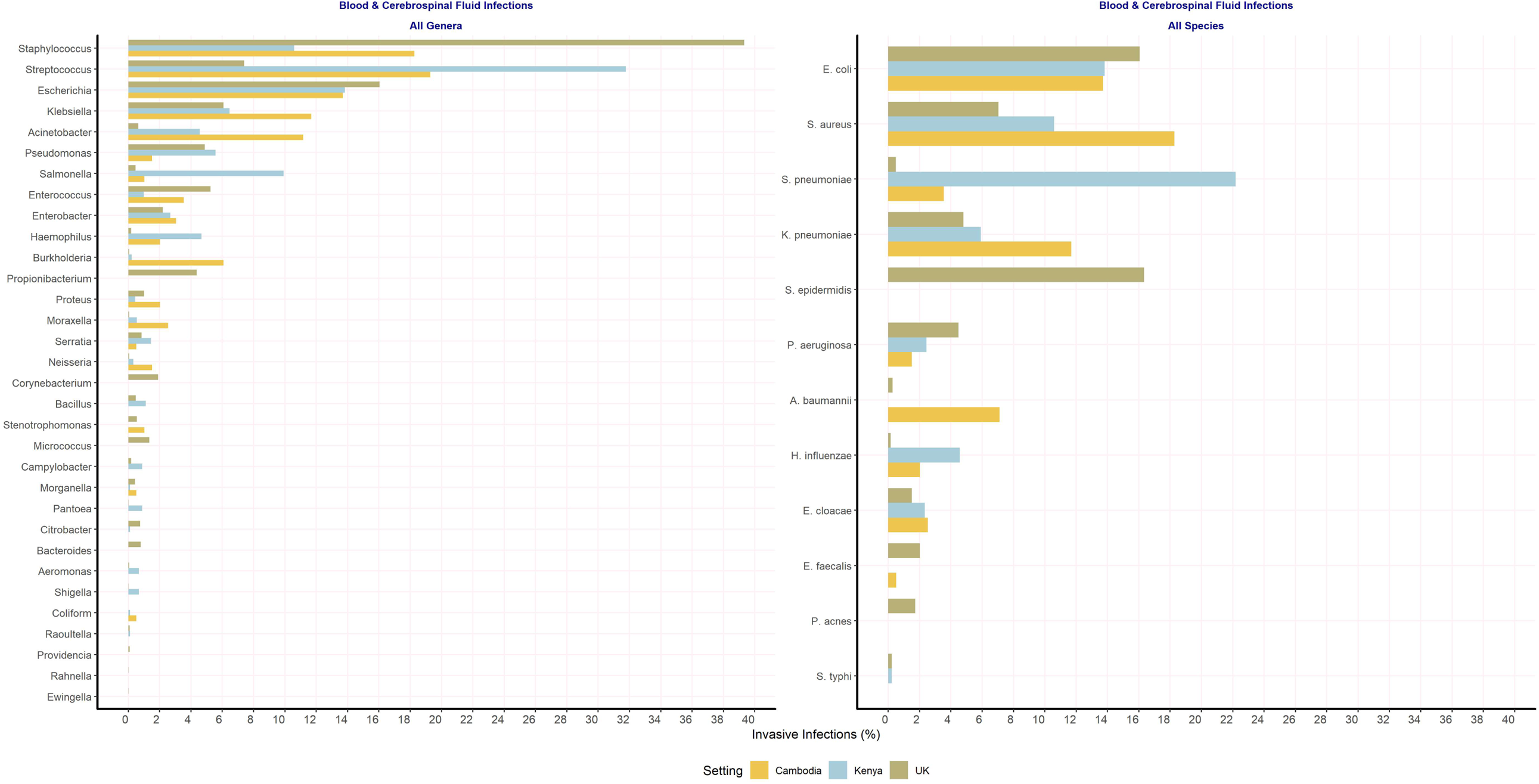

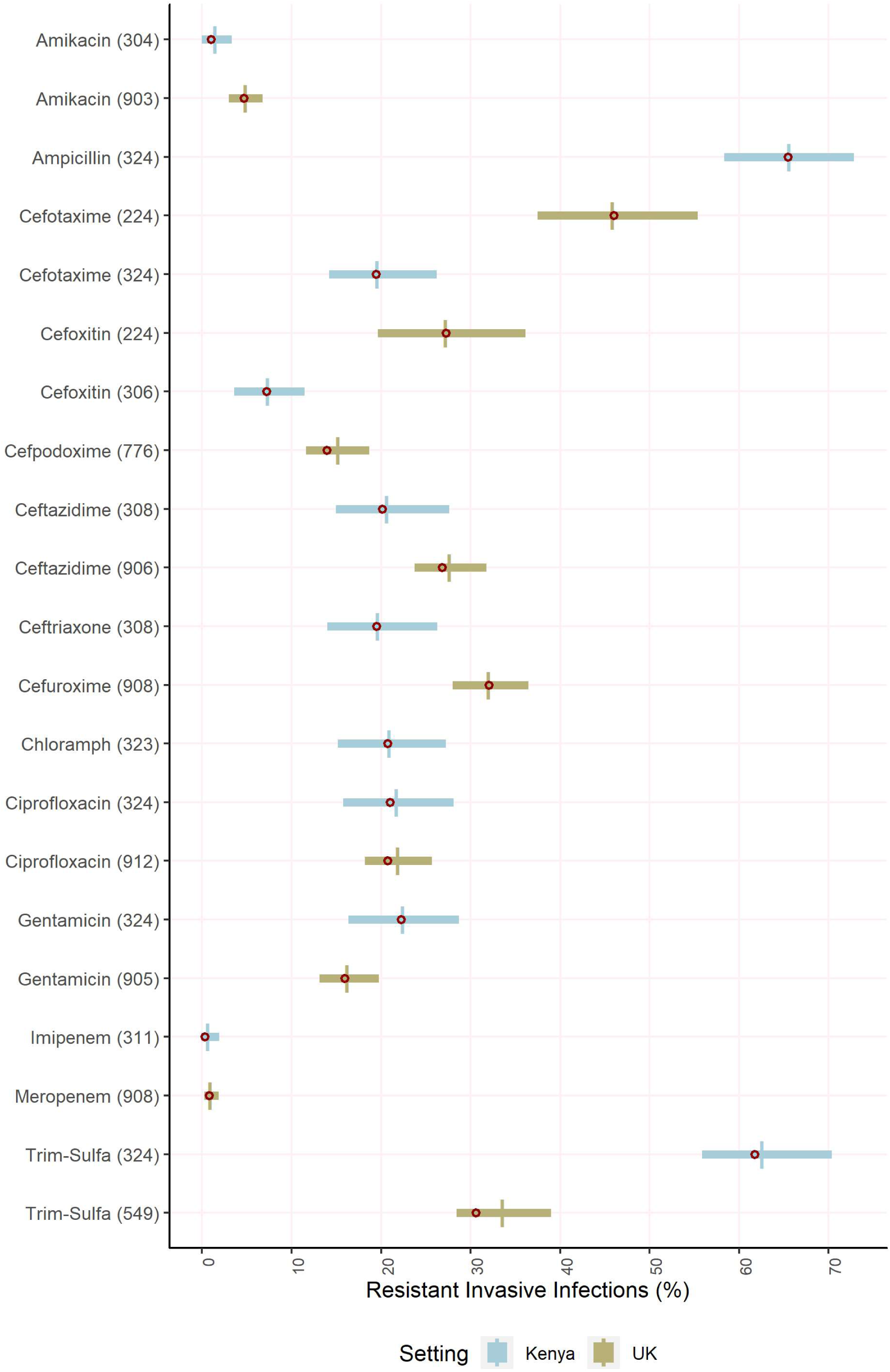

