## Supplementary Methods for "Metrics for Public Health Perspective Surveillance of Bacterial Antibiotic Resistance in Low- and Middle-Income Countries"

### Supplementary Methods 1.

#### Modified MoBio PowerSoil DNA Extraction Method for Metagenomic DNA Extraction from Rectal Swabs

Rectal swabs suspended in Phosphate Buffer Saline (PBS) or Amies trasport media (ATM) and stored at -80°C, were thawed on ice. The MoBio PowerSoil DNA Isolation Kit (Qiagen, Hilden, Germany) was used for DNA extraction using the following modified SOP:

##### a) Sample Preparation and Initial Lysing Steps

Before getting started: Check Solution C1. If Solution C1 is precipitated, heat solution to 60°C until dissolved before use.

1. Transfer swab head into the PowerBead tube without vortexing the thawed sample.
2. Centrifuge the original sample tube, which now only contains PBS or ATM, at 13,200 rpm for 10 min.
3. Carefully remove the supernatant from the original sample tube (about 800ul of PBS or ATM) without disturbing the pellet.
4. Re-suspend the pellet using the remaining 200 µl of PBS or ATM in the sample tube.
5. Transfer the mix into the PowerBead tube.
6. Add *Thermus thermophilus*[1] DNA (8.75 ul/sample [1ng/ul]).
7. Add 60 µl of Solution C1 to the PowerBead tube.
8. Vortex for 5 seconds.
9. Bead beat[2] PowerBead tubes using the following set up: 6.0 m/s, 40 seconds, 2 cycles, 300 second pause, 1 ml volume, Lysing matrix E. Incubate rack and tubes at 4°C between cycles.

##### b) DNA extraction

1. Incubate PowerBead tubes at 65°C for 10 minutes.
2. Vortex for 5 seconds.
3. Incubate PowerBead tubes at 65°C for another 10 minutes.
4. Vortex for 5 seconds.
5. Incubate PowerBead tubes at 95°C for 10 minutes.
6. Incubate PowerBead tubes at 4°C for 5 minutes.
7. Bead beat[2] PowerBead Tubes using the following set up: 6.0 m/s, 40 seconds, 2 cycles, 300 second pause, 1 ml volume, Lysing matrix E. Incubate rack and tubes at 4°C between cycles.

---

[1] NB - this step is not absolutely essential. This is used in our metagenomics workflows for individual samples, where it is of particular value in the normalisation of gene counts (please see Gweon et al, Environmental Microbiome volume 14, Article number: 7 (2019), https://environmentalmicrobiome.biomedcentral.com/articles/10.1186/s40793-019-0347-1).

[2] Rectal Swabs in PBS were processed at the Nuffield Department of Medicine (University of Oxford, UK) using the FastPrep-24 5G instrument (MP Biomedicals, Santa Ana, CA, USA). Rectal swabs in ATM were processed at the Clinical Infection and Diagnostics Research laboratory at Guy's and St Thomas' NHS Foundation Trust, UK. A FastPrep instrument or equivalent was not available in the latter. Bead beating steps were hence replaced by vortexing at maximum speed on the Labnet VX-100 vortex for 30 minutes.

8. Centrifuge tubes at 10,000 x g for 30 seconds at room temperature. Make sure the PowerBead tubes rotate freely in your centrifuge without rubbing.

CAUTION: Be sure not to exceed 10,000 x g or tubes may break.

9. Transfer 600 µl of the supernatant to a clean 2 ml Collection Tube.
10. Add 250 µl of Solution C2.
11. Vortex for 5 seconds.
12. Incubate at 4°C for 5 minutes.
13. Centrifuge the tubes at room temperature for 2 minutes at 10,000 x g.
14. Avoiding the pellet, transfer up to, but no more than, 600 µl of supernatant to a clean 2 ml Collection Tube.
15. Add 200 µl of Solution C3.
16. Vortex briefly.
17. Incubate at 4°C for 5 minutes.
18. Centrifuge the tubes at room temperature for 1 minute at 10,000 x g.
19. Avoiding the pellet, transfer up to, but no more than, 750 µl of supernatant into a clean 2 ml Collection Tube.
20. Shake to mix Solution C4 before use.
21. Add 1200 µl of Solution C4 to the supernatant and vortex for 5 seconds.
22. Load approximately 675 µl onto a Spin Filter and centrifuge at 10,000x g for 1 minute at room temperature. Discard the flow through and add an additional 675 µl of supernatant to the Spin Filter and centrifuge at 10,000 x g for 1 minute at room temperature. Load the remaining supernatant onto the Spin Filter and centrifuge at 10,000 x g for 1 minute at room temperature.

NOTE: A total of three loads for each sample processed are required.

23. Add 500 µl of Solution C5 and centrifuge at room temperature for 30 seconds at 10,000 x g.
24. Discard the flow through.
25. Centrifuge again at room temperature for 1 minute at 10,000 x g.
26. Carefully place spin filter in a clean 2 ml Collection Tube. Avoid splashing any Solution C5 onto the Spin Filter.
27. Add 50 µl of Solution C6 to the centre of the white filter membrane.
28. Incubate Spin Filters at room temperature for 2 minutes.
29. Centrifuge at room temperature for 30 seconds at 10,000 x g.
30. Discard the Spin Filter. The DNA in the tube is now ready for any downstream application. No further steps are required.

### Supplementary Methods 2.

#### Modified MoBio PowerSoil DNA Extraction Method for Metagenomic DNA Extraction from faecal slurry samples

Faecal slurry samples (faeces suspended in nutrient broth +10% glycerol) stored at -80°C, were thawed on ice. The MoBio PowerSoil DNA Isolation Kit (Qiagen., Hilden, Germany) was used for DNA extraction using the following modified SOP:

##### a) Assess Pellet size

1. Weigh and record empty microcentrifuge tube weights
2. Vortex thawed samples and transfer samples to pre-weighed microcentrifuge tubes using wide orifice pipette tips
3. Centrifuge at 13,200 rpm for 10 min
4. Carefully remove the supernatant without disturbing the pellet
5. Re-weigh microcentrifuge tube and calculate pellet weight
6. Resuspend the pellet using 500 µl of buffer from a PowerSoil® DNA Isolation Kit Bead tube using wide orifice pipette tips
7. Transfer the mix back into the Bead tube and proceed with DNA extraction or store at -80°C for future extraction

##### b) DNA extraction

Before getting started: Check Solution C1. If Solution C1 is precipitated, heat solution to 60°C until dissolved before use.

8. Add *Thermus thermophilus*[3] DNA (8.75 ul/sample [1ng/ul])
9. Vortex the PowerBead tube containing resuspended faecal pellet for 3 seconds.
10. Add 60 µl of Solution C1 and vortex for 5 seconds.
11. Incubate PowerBead Tubes at 65°C for 10 minutes.
12. Vortex for 5 seconds.
13. Incubate PowerBead Tubes at 65°C for another 10 minutes.
14. Vortex 5 seconds.
15. Incubate PowerBead Tubes at 95°C for 10 minutes.
16. Incubate PowerBead Tubes at 4°C for 5 minutes.
17. Bead Beat PowerBead Tubes using the following protocol: 6.0 m/s, 40 seconds, 2 cycles, 300 second pause, 1 ml volume, Lysing matrix E. Incubate rack and tubes at 4°C between cycles.

[3] NB - this step is not absolutely essential. This is used in our metagenomics workflows for individual samples, where it is of particular value in the normalisation of gene counts (please see Gweon et al, Environmental Microbiome volume 14, Article number: 7 (2019), https://environmentalmicrobiome.biomedcentral.com/articles/10.1186/s40793-019-0347-1).

18. Make sure the PowerBead Tubes rotate freely in your centrifuge without rubbing. Centrifuge tubes at 10,000 x g for 30 seconds at room temperature.

CAUTION: Be sure not to exceed 10,000 x g or tubes may break.

19. Transfer 600 µl of the supernatant to a clean 2 ml Collection Tube.
20. Add 250 µl of Solution C2 and vortex for 5 seconds. Incubate at 4°C for 5 minutes.
21. Centrifuge the tubes at room temperature for 2 minutes at 10,000 x g.
22. Avoiding the pellet, transfer up to, but no more than, 600 µl of supernatant to a clean 2 ml Collection Tube.
23. Add 200 µl of Solution C3 and vortex briefly. Incubate at 4°C for 5 minutes.
24. Centrifuge the tubes at room temperature for 1 minute at 10,000 x g.
25. Avoiding the pellet, transfer up to, but no more than, 750 µl of supernatant into a clean 2 ml Collection Tube.
26. Shake to mix Solution C4 before use. Add 1200 µl of Solution C4 to the supernatant and vortex for 5 seconds.
27. Load approximately 675 µl onto a Spin Filter and centrifuge at 10,000x g for 1 minute at room temperature. Discard the flow through and add an additional 675 µl of supernatant to the Spin Filter and centrifuge at 10,000 x g for 1 minute at room temperature. Load the remaining supernatant onto the Spin Filter and centrifuge at 10,000 x g for 1 minute at room temperature. A total of three loads for each sample processed are required.
28. Add 500 µl of Solution C5 and centrifuge at room temperature for 30 seconds at 10,000 x g.
29. Discard the flow through.
30. Centrifuge again at room temperature for 1 minute at 10,000 x g.
31. Carefully place spin filter in a clean 2 ml Collection Tube. Avoid splashing any Solution C5 onto the Spin Filter.
32. Add 70 µl of Solution C6 to the centre of the white filter membrane.
33. Incubate Spin Filters at room temperature for 2 minutes.
34. Centrifuge at room temperature for 30 seconds at 10,000 x g.
35. Discard the Spin Filter. The DNA in the tube is now ready for any downstream application. No further steps are required.
