## Supplementary Figures for "Metrics for Public Health Perspective Surveillance of Bacterial Antibiotic Resistance in Low- and Middle-Income Countries"

### Supplementary Fig 1

**Main species and genera (all bacterial orders) identified from blood and cerebrospinal fluid infections in Cambodia, Kenya and UK.**

Panels show, for each setting, percentages of the most common bacterial species and genera out of all bacterial infection isolates with speciation results identified from blood and cerebrospinal fluid samples in target age groups, from 2010-2017 (Cambodia [n=197]; Kenya [n=910]; UK [n=3356]).

### Supplementary Fig 2

**Bayesian model prediction of proportion of Enterobacterales invasive infections with resistance to antibiotics. Predictions shown for antibiotics where antibiotic susceptibility test (AST) results were available from > 100 invasive infection isolates.**

Horizontal bars represent 95% highest density posterior interval and vertical lines represent means of the model predictions based on metagenomic data from population pools. Red circles show the proportion of blood and cerebrospinal fluid Enterobacterales infections that were found to be resistant to the antibiotic listed in the y-axis. The number of isolates with AST results are also given in the y-axis. Percentages were calculated by dividing observed and predicted counts by the total number of invasive infection isolates with AST data in each setting (see y-axis). “Trimethoprim.” is short for trimethoprim-sulfamethoxazole; “Cloramph” is short for chloramphenicol.
